# It takes neurons to understand neurons: Digital twins of visual cortex synthesize neural metamers

**DOI:** 10.1101/2022.12.09.519708

**Authors:** Erick Cobos, Taliah Muhammad, Paul G. Fahey, Zhiwei Ding, Zhuokun Ding, Jacob Reimer, Fabian H. Sinz, Andreas S. Tolias

## Abstract

Metamers, images that are perceived as equal, are a useful tool to study representations of natural images in biological and artificial vision systems. We synthesized metamers for the mouse visual system by inverting a deep encoding model to find an image that matched the observed neural activity to the original presented image. When testing the resulting images in physiological experiments we found that they most closely reproduced the neural activity of the original image when compared to other decoding methods, even when tested in a different animal whose neural activity was not used to produce the metamer. This demonstrates that deep encoding models do capture general characteristic properties of biological visual systems and can be used to define a meaningful perceptual loss for the visual system.

## Main

To decipher the brain’s algorithms of perception it is critical to understand how natural visual scenes are represented in neural activity. This question has traditionally been studied with two complementary approaches: *encoding* methods characterize neural activations as a function of the stimulus, while *decoding* methods reconstruct the visual input or specific stimulus features from neuronal recordings. The ability to record large scale neuronal activity in combination with recent advances in machine learning have substantially increased the accuracy of encoding and decoding models including in higher visual areas [1–5]. While encoding models have a clear quality metric—the prediction accuracy of neural activity—interpreting them can be challenging when modeling responses to natural images given that neuronal tuning is highly nonlinear [1, 3]. Decoding models, on the other hand, can yield interpretable stimulus variables represented in specific neuronal populations such as stimulus orientation, the direction of motion of a stimulus or even entire scenes [4, 6–15]. In particular, decoding models can be used to produce “metamers”, stimuli that are perceived as the same or produce the same activity in a set of units in biological or artificial networks [16, 17].

However, decoding algorithms are fundamentally determined by the choice of quality metric, i.e. training loss function, used to estimate the decoded variables. In the case of decoding specific latent variables such as the class of an object, the loss function is clearly defined. In more challenging problems like reconstructing entire visual scenes, the loss function is less clear. Image-based metrics, such as the commonly used mean-squared-error of pixel intensities, are not well correlated with perceptual similarity [18]. Moreover, devising a good perceptual loss function for reconstructing complex natural scenes is especially difficult for higher visual areas, specialized on representing specific latent variables from visual scenes such as textures, shapes, colors and faces. Ultimately, what matters most is how well the decoded stimuli reproduce the original brain activity used to reconstruct them, i.e., synthesizing metamers of the original image with respect to a neuronal populations. Unfortunately, experimentally it is practically impossible to decode complex stimuli – such as natural scenes – based on a comparison between the neural activity elicited by the original and the decoded image recorded in real time. Here, we reconstructed natural images by “inverting” a deep learning based encoding model (Fig. 1). The objetive of the inversion process was to synthesize stimuli such that the activity predicted by the encoding model matched the recorded brain activity to the original image. Measuring the reconstruction quality on the neural responses, allowed us to obtain stimuli that were neurally equivalent to the original natural image. When we showed these reconstructed images back to the animals they more accurately reproduced the original brain activity than images decoded from other algorithms we tested, even when shown to different neural populations or even different animals.

**Figure 1.**
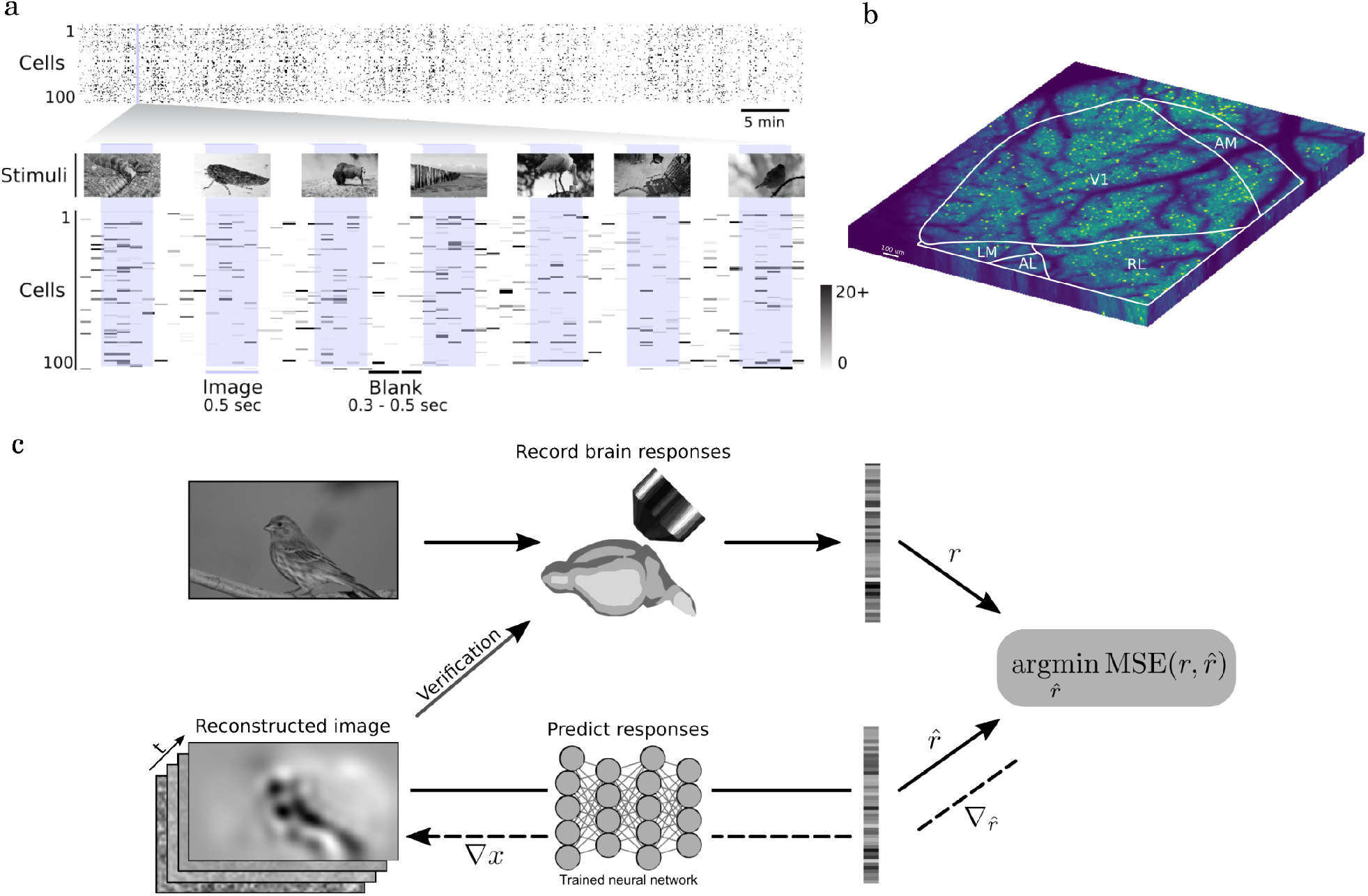
Data recording paradigm. **a)** Our stimuli for training the deep encoding model are composed of 5040 natural images shown for 0.5 seconds interleaved with blank spaces (0.3-0.5 seconds). **b)** We record approximately 8000 cells across four visual areas in a 2.4 × 1.8 *mm*^2^ field of view. **c)** Schematic of our proposed reconstruction method. We train a state-of-the-art differentiable encoding model to predict brain responses and use it to reconstruct arbitrary images not previously shown to the model. To reconstruct images, we start with a blank image and iteratively modify it to minimize the distance between the in-vivo recorded response and the prediction from the encoding model using backpropagation. To evaluate the proposed method we later show the reconstructions, along with the original images to a similar population of cells in visual cortex.

We recorded the responses of thousands of excitatory neurons in layer 2/3 across multiple areas of the mouse visual cortex using two-photon imaging with a wide field-of-view mesoscope to natural images (Fig 1) [19]. Next, we trained a deep convolutional neural network (CNN) to predict neural responses as a function of the visual input and measured correlates of behavioral modulation including running, eye movements and pupil dilation. The encoding model was trained on 5000 natural images and accurately predicted the activity of thousands of neurons recorded from different cortical visual areas on a held out test set of natural images. On average across scans, we obtained a normalized correlation coefficient of 0.61 (absolute correlation coefficient of 0.56) [20]. This deep neural network encoding model provides an accurate digital twin of the mouse visual cortex and was previously used to find most exciting input for single neurons [1]. To decode from the digital twin an initially blank image was optimized via gradient descent to produce predicted responses that matched the *in vivo* recorded responses while softly smoothing the image after each iteration to avoid unrealistic high-frequency details (Fig 1c). For comparison, we also decoded images using three algorithms optimized to directly reproduce the presented images in pixel space: an L2-regularized linear regression (Linear), a two-layer multi-layer perceptron (MLP) and a deep neural network with transposed convolutions (Deconv) [21]. Moreover, we decoded from a Gabor filter bank encoding model of visual cortex (Gabor) [5] and a model that samples natural images to create reconstructions (AHP) [4]; see Methods for further details. Because of the large number of neurons (approx. 8200 per mouse), all decoding methods were able to produce reconstructions that resembled the original image (Fig. 2a, 3a)

**Figure 2.**
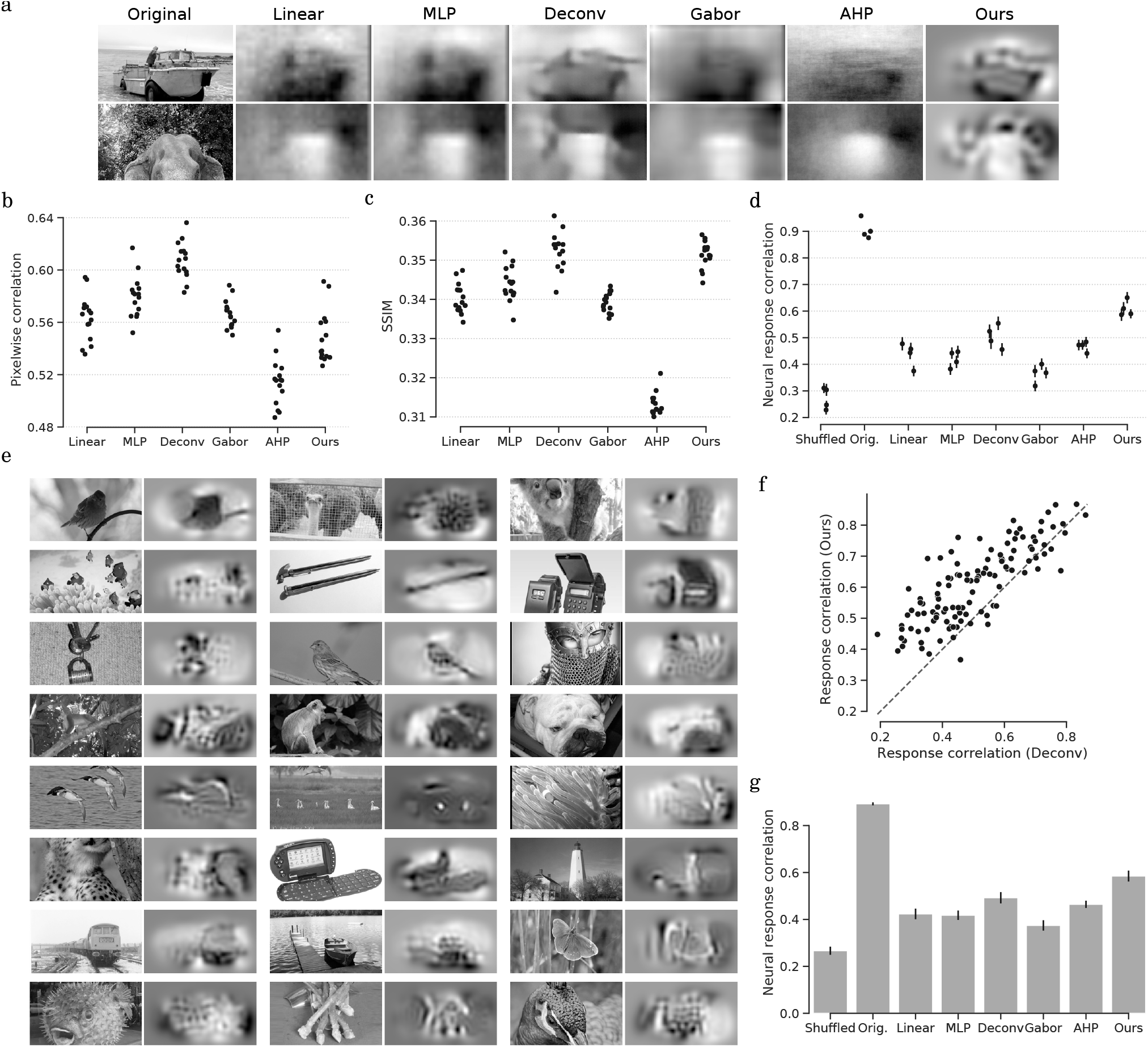
Evaluation of natural image reconstructions. **a)** Sample reconstructions for the six tested methods (See Sup. Fig. S1 for other examples.) **b, c)** Swarm plots (*n* = 15 scans) of the pixel-wise correlation (b) and structural similarity (c) between original images and their reconstructions (averaged across all images). **d)** Correlation between neural response to the original image and to the reconstructed image (*n* = 4 scans). Each dot shows the mean correlation across images for one scan (error bar is the standard error of the mean). ‘Shuffled’ is the expected correlation between responses to two unrelated natural images estimated as the average of the off-diagonal values in the image-to-image response correlation matrix while ‘Orig.’ is the correlation of neural responses to two distinct presentations of the same image (average of the diagonal of the correlation matrix); these provide a lower and upper bound for the correlation between images and reconstructions in a scan. **e)** Sample reconstructions from our method. The windowing effect results from the field of view of recorded cells not covering the monitor fully. **f)** Correlation of neural responses (*n* = 4 scans) between original images and reconstructions from a deconvolution network (x axis) and from our method (y axis). In-vivo responses elicited by our method are more similar to those of the original image (0.61 vs 0.51 mean correlation, t-test p-value=1e-8). **g)** Correlation between neural responses (*n* = 1 scan) from the original images and reconstructions generated for a different mouse. Error bars show standard error across images.

**Figure 3.**
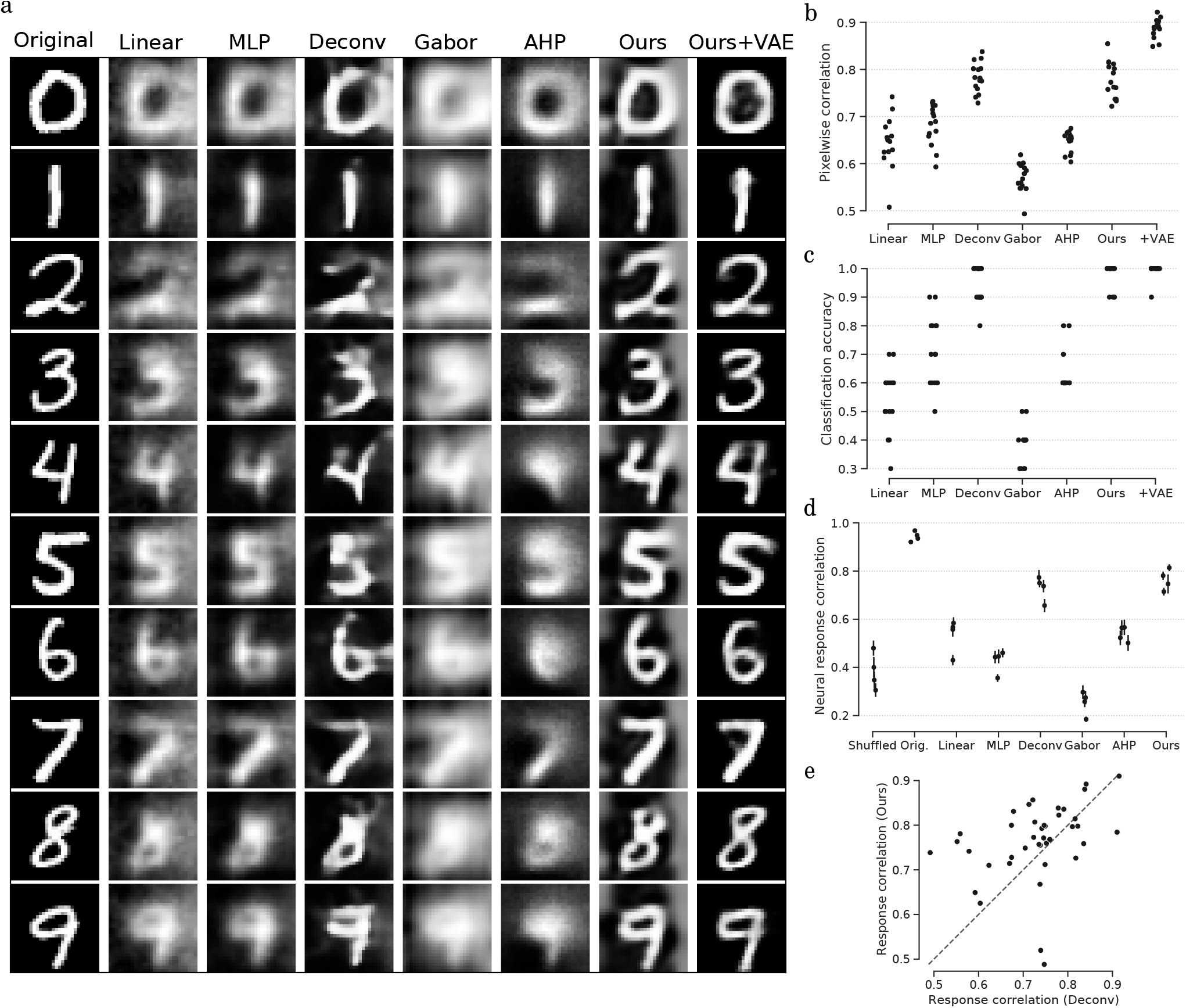
Evaluation for MNIST images. **a)** MNIST reconstructions for seven different methods. **b)** Swarm plots (*n* = 15 scans) of pixel-wise correlation between original images and their reconstructions (averaged across all images) for the different methods. **c)** Swarm plots (*n* = 15 scans) for the accuracy of a pretrained MNIST classifier on the reconstructions from different methods. **d)** Correlation between the neural response to the original digit and the reconstruction. Each dot represents the average correlation (*n* = 4 scans) and the error bars show the standard error of the mean. ‘Shuffled’ is the expected correlation between the neural responses to two unrelated digit images estimated as the average of the off-diagonal values in the image-to-image correlation matrix while ‘Orig.’ is the correlation between neural responses to two distinct presentations of the same image (average of the diagonal of the correlation matrix); these are a lower and upper bound for the correlation between images and reconstructions. **e)** Correlation of neural responses (*n* = 4 scans) between original images and the reconstructions from the deconvolution method (x axis) and from our method (y axis). Neural responses elicited by our method are not significantly different than those from the deconvolution method (0.76 vs 0.73 mean correlation, t-test p-value=0.09)

We compared the different reconstructions using three metrics: 1) correlation between pixel intensities, 2) the structural similarity index (SSIM) [22]—a widely used measure to judge the perceptual quality of an image compared to a reference image, and 3) the correlation of neural activity between the original image and the one elicited by its reconstruction in subsequent *in vivo* closed-loop experiments. Specifically, we presented the original images and the reconstructions to the same or a different mouse in the same visual areas and cortical layer but not necessarily to the same neurons. This is a strong test assessing whether the reconstruction method puts emphasis on image features that matter for cortex. To the best of our knowledge, this is the first time that the quality of decoded stimuli is quantified in physiological experiments.

As expected, on the pixelwise corrrelation metric, reconstruction methods optimized to directly reproduce the presented images pixel-by-pixel (e.g., MLPs or deep deconvolution networks) outperformed the others including our method of inverting deep learning encoding models (Fig. 2b, Sup. Tab. 1). On the SSIM metric our method was on par with the pixel deconvolutional network method (Deconv, Fig. 2c). Our method produced more perceptually meaningful reconstructions (Fig. 2e, Sup. Fig. S1) and yielded the best match between the neural activity elicited between the reconstructed and original image (Fig. 2d, f, Sup. Tab. 1). Importantly, our encoding based reconstructions also generalized better across animals (Fig. 2g). Therefore, these closed loop experiments highlight that image-based decoding methods and quality metrics such as pixel level training loss emphasize unimportant features of decoded stimuli. In contrast, our method of inverting accurate digital twin models of brain activity synthesize metamers of the original images with respect to brain activity emphasizing meaningful features for specific neuronal populations – even across animals.

We further evaluated the decoding methods in a dataset of grayscale images of handwritten digits [23] akin to datasets used in earlier decoding research [24, 25]. Even though the deep encoding model was trained on responses to natural images only, our approach produced quantitatively and qualitatively better reconstructions (Fig. 3). Additionally, we prepended an image generation model (a pretrained variational auto encoder VAE [26]) to our model and performed gradient descent in the latent image representation of this generator rather than on the pixels. This method reconstructed images that were almost identical to the original input using only *in vivo* neural responses (Ours+VAE in Fig. 3a).

Our results suggest that deep neural encoding models trained on the responses to natural stimuli can be used to define an image similarity metric that produces metamers that best match neural activity, across neurons and animals. These images yield visually accessible qualitative features that are represented in different parts of the visual systems, and can be quantitatively verified in physiological experiments.

## Methods

### Experimental setup

All procedures were approved by the Institutional Animal Care and Use Committee (IACUC) of Baylor College of Medicine. We ran our experiments in seven mice (*Mus musculus*, 4 female, 3 male) aged 70-188 days old (129 on average) expressing GCaMP6s in excitatory neurons via SLC17a7-Cre and Ai162 transgenic lines (JAX stock #023527 and #031562, respectively). We made a 4 mm diameter craniotomy over visual cortex as previously described [27].

During experiments, mice were head-mounted above a cylindrical treadmill and allowed to run freely. Calcium imaging was performed using a Chameleon Ti-Sapphire laser (Coherent) tuned to 920 nm and a large field of view mesoscope [19] with a custom objective (0.6 NA, 21mm focal length); laser power after the objective was kept below 60 mW to avoid tissue damage. We presented visual stimuli to the left eye with a 25” LCD monitor (ASUS PB258Q, 1920 × 1080 px resolution) positioned 15 centimeters away from the eye. Rostro-caudal treadmill movement was measured using a rotary optical encoder with a resolution of 8000 pulses per revolution. To capture eye movements, light diffusing from the laser through the pupil was reflected through a hot mirror and captured with GigE CMOS camera (Genie Nano C1920M, Teledyne Dalsa) at 20 fps at 1920 × 1200 px resolution.

### Image stimulus

The stimulus consists of 6200 images: 5000 natural images shown once, 30 natural images shown 40 times each and 10 MNIST digit images shown 40 times. Natural images were sampled at random from the ImageNet dataset [28]: ImageNet images have arbitrary dimensions, we take the maximal 16:9 ratio crop from the center of the image, resize it to 144 × 256 and turn it into grayscale. From an initial random sample of 100 images, we hand-selected 30 natural images—the ones shown for more repetitions—to make sure they show interesting structures; this selection was done before any experiments. The 10 MNIST images [23] were chosen manually from the standard test set to have the canonical appearance of each digit. MNIST images are 28 × 28 pixels: we resized them to 144 × 144, rotate them 90 degrees and pad them with black pixels to fill the 144 × 256; we assume that most digits are thin and tall so we rotate them to lessen effects from the edge of the monitor atop and at the bottom of the digit.

During stimulus presentation, images are interleaved and shown for 0.5 seconds with a blanking period—where the screen is shown in gray—of 300-500 milliseconds between them. The length of the blanking period is chosen uniformly at random.

### Imaging

Before experiments, we recorded pixel-wise responses (at 0.2 px/*µ*m) to drifting bar stimuli for a 2.4 × 2.4 *mm*^2^ region of interest at 200 *µ*m depth from the cortical surface to generate a sign map for delineating visual areas [29]. We chose an imaging site to maximally cover visual areas while avoiding blood vessel occlusion and leveled the craniotomy window to keep the surface of the brain parallel to the objective using the scan’s six degrees of freedom.

During experiments, using a remote objective, we sequentially image three 2400 × 620 *µ*m^2^ fields per frame at 8.1 fps and 0.4 px/*µ*m xy-resolution. Fields are recorded side by side with 20um overlap to cover a 2.4 × 1.8*mm*^2^ recording plane in layer 2/3 at around 200 *µ*m depth from the surface of the cortex.

### Processing of neural and behavioral data

Each frame of the recording field is motion corrected using rigid template matching; the template is obtained by averaging 2000 frames (approx 4 minutes) from the middle of the scan. Cell masks and fluorescence traces are obtained using constrained non-negative matrix factorization [30] and spike activity is deconvolved from these fluorescence traces using the same package. For further analysis, we restrict to cells in visual areas (V1, LM, AL and RL) that are at least 8 microns from the edge of any recording field (this avoids any edge effects from recording).This results in approximately 8200 cells per scan on average. We subsequently average the spike traces over the 500 ms duration of each image (adding 30 ms start offset meant to account for the time the signal takes to travel from the retina to cortex [31]) to obtain a single response vector per image.

The encoding model receives treadmill velocity, pupil size, and pupil location as auxiliary signals during training. To obtain treadmill velocity, we take the numerical gradient of the wheel location using central differences [32]. The pupil is segmented from each video camera frame using DeepLabCut [33], a circle is fitted to the predicted mask and its diameter and position is used as pupil size and location. These behavioral traces are also averaged during image presentation to obtain a single number per image. We used the open-source DataJoint framework for the computational workflows [34, 35].

### Closed loop

To evaluate reconstructions in neural space, we are able to show the reconstructions from one day back to the mice on the next day. To avoid long calcium imaging sessions, we evaluate the reconstructions from three methods at a time in a single scan ^1^: we create a stimulus with reconstructions for the 40 test images (30 natural + 10 digits) repeated 40 times and the original images repeated 60 times (7200 images in total). Stimulus presentation, recording and data processing parameters are the same as for the original experiments. We recorded five closed loop scans in total.

From one day to the next, all but the z axis of the scan were locked to allow us to return to the same imaging site. Although we do not perform day to day cell matching, field of views look visually similar and we expect the neural population across days to overlap highly. In order to have a consistent monitor placement relative to the mouse, we placed the aggregate receptive field for each day at the center of the monitor. To obtain the aggregate receptive field we tiled the center of the screen in a 10 × 10 grid with single dark dots over bright background (∼5^°^) and averaged the calcium trace of a ∼150 × 150*µ*m^2^ window in the center of our field-of-view from 0.5–1.5 seconds after stimulus onset across all repetitions of the stimulus for each location. We fitted a ellipsis to this coarse grid and displaced the monitor each day to keep the field-of-view centered.

## Machine Learning model

### Data

We train the models using 4500 natural images and their recorded neural responses and select hyper- parameters with the remaining 500 images. The 30 natural images with repeats (see Sec.) are used for evaluation: we average the neural responses across repeats to increase signal-to-noise ratio. We isotropically downsampled stimuli images to 36 × 64 pixels. Input images, the target neuronal activities, behavioral traces, and pupil positions were normalized using the mean and standard deviation from the training set.

### Reconstruction based on deep encoding models

Rather than training a decoding model to predict images from neural responses, we train a differentiable encoding model and pseudo-invert it using its gradients to obtain a predicted reconstruction.

The encoding model is a three layer neural network that extracts intermediate image features and a readout layer that learns the position of each cell’s receptive field in the monitor, extracts the intermediate features at that point and linearly predicts a cell response [1, 36]. To obtain reconstructions, we feed a blank initial image to the encoding model and iteratively optimize it via gradient descent so that the predicted neuronal response matches the recorded neuronal response [37]. We use an automatic differentiation engine [38] to compute the gradient of the mean squared error between predicted and recorded neural responses with respect to the input image. We gaussian blur the gradient image (standard deviation = 2.5px) each iteration to avoid high frequency noise in the reconstruction [39]. We run the optimization for 1000 steps: it takes around 5 secs in a modern machine. Final reconstructions are bilinearly upsampled to their original size (144 × 256) and the contrast is scaled to cover the full range of intensity values (0-255).

Minimizing distances between neural responses is more sensible than minimizing distances in the high dimensional image space used in standard decoding models. The inductive biases learned by encoding models for the simpler task of neural response prediction may also result in the better reconstructions produced by our method. Another advantage of our reconstruction method is that we can use a (possibly hand-crafted or pre-trained) generative model to create the images fed to the encoding model and optimize directly on the latent space of the generative model rather than in the space of images; this adds a heavy natural bias on the type of reconstructions we are able to produce and allows us to take advantage of powerful generative models from the deep learning literature [40–42]. We show this using a variational autoencoder pre-trained on the MNIST digit images [26]: we optimize on the 20-dimensional latent space of the autoencoder via gradient descent without any gaussian blurring. Reconstructions look strikingly similar to the original images (see Ours+VAE in Fig. 3a)

### Evaluation

Evaluating the quality of natural image reconstructions is not straightforward and the metrics used may skew results. We report the two most commonly used similarity metrics: pixel-level correlation and structural similarity. For any hyperparameter selection we prefer to use structural similarity as, from experience, it aligns better with the human notion of a good reconstruction and is less biased towards preferring blurry reconstructions. For reconstruction of MNIST images, we use the performance of an MNIST classifier on the reconstructed image as an additional measure of reconstruction quality. We measure this with a pre-trained three layer MLP with an accuracy of 98% on the original MNIST test set.

In a separate experiment, we show the reconstructions back to the mice along with the original images (see Sec.); this allows us to compute the similarity of neural responses in the brain, arguably the true objective of decoding methods. We average recorded responses across trials to compute the correlation between responses to original images and reconstructions. Because of neuronal variability and instrument noise, response correlations will never be perfect even when presenting the same image. To estimate an upper bound on the possible response correlation in a scan, we split the 60 image trials into two disjoint subsets of 30 trials and compute the correlation between them (averaged across images). We also estimate a lower bound on the response correlation by computing the correlation of responses to different images (averaged across all pairs of distinct images); non-zero correlations arise from natural neuron-to-neuron covariance and image similarities.

### Comparisons to other decoding methods

We compare six different methods. We search across different hyperparameters for each to obtain the best performing model (see Sup. Sec. A.5).

- Linear: A linear regression model from responses to pixels regularized with either an L2 or L1 penalty. Hyperparameters: image dimensions and regularization weight.
- MLP: A multilayer perceptron with a single hidden layer. Hyperparameters: image dimensions, learning rate, L2 regularization weight and number of units in the hidden layer.
- Deconvolution network (Deconv): An eleven layer convolutional network with transposed convolution layers for upsampling. The initial layer is a transposed convolution that upsamples the 1 × 1 neural response to 3 × 5, afterwards every second convolution upsamples its input by a factor of two. Hyperparameters: learning rate and L2 regularization weight.
- Gabor [5]: Filter all images using a pre-defined Gabor filter bank, learn to predict the weights for these filters using linear regression (with either L1 or L2 regularization) and create reconstructions by linearly combining the Gabor filters with the predicted weights. Current state of the art in mice reconstruction. Hyperparameters: image dimensions, size of the Gabor filter bank and regularization weight.
- Averaged high posterior (AHP) [4]: Pass many natural images through the encoding model and average the images that produce a closer predicted response to the target neural response. We use a bank of 125,000 images that were not used in any other experiment. Hyperparameters: number of images to sample, whether or not to use a weighted average of images (using the normalized similarity value between responses as weights).
- Ours: As explained in the previous sections. We treat the optimization step size and sigma for gaussian blurring as hyperparameters. Number of iterations used is chosen manually to be large enough that the optimization converges (1000 in our case).
- Ours + VAE: As explained in the previous sections. No hyperparameters; we manually searched for a sensible step size and used it for all scans.

## Acknowledgements

The authors thank Stelios Papadopoulos and the members of the Tolias lab for productive discussions and feedback on the project.

This work was supported by the Intelligence Advanced Research Projects Activity (IARPA) via Department of Interior/Interior Business Center (DoI/IBC) contract number D16PC00003. The U.S. Government is authorized to reproduce and distribute reprints for Governmental purposes notwithstanding any copyright annotation thereon. Disclaimer: The views and conclusions contained herein are those of the authors and should not be interpreted as necessarily representing the official policies or endorsements, either expressed or implied, of IARPA, DoI/IBC, or the U.S. Government. The project was also supported by DARPA grants 5N66001 and N6001-19-C-4020 and 5N66001 from the NESD and MOANA programs respectively. FHS was supported by the Carl-Zeiss-Stiftung and acknowledges the support of the DFG Cluster of Excellence “Machine Learning – New Perspectives for Science”, EXC 2064/1, project number 390727645.

## Author contributions statement

T.M. and P.G.F. recorded the scans. Z.D and Z.D processed the scans to obtain the encoding datasets. J.R. provided supervision and technical input regarding recording. A.S.T., F.H.S. conceptualized the experiments, provided supervision and wrote the article. E.C implemented the processing pipeline for scans, implemented the decoding methods, analyzed the results and wrote the article. All authors reviewed the manuscript.

## A Supplementary section

### A.1 Quantitative evaluation of results

We report results using three metrics: 1) pixel-wise correlation between original image and reconstruction, 2) structural similarity between original image and reconstruction and 3) correlation of responses elicited in the brain when shown original and reconstructed image. We estimate an upper bound on the brain response correlation by computing the correlation between responses to presentations of the same image: we average the responses to two sets of 30 distinct presentations of each natural image and compute the correlation using those averages. The upper bound on the neuronal response correlation for natural images was 0.908; the remaining gap to perfect image-to-image correlation can be attributed to natural response variability and recording limitations.

**Table 1.** Evaluation metrics for natural images averaged across scans.

| Method | Pixel-wise correlation | SSIM | Response correlation |
| --- | --- | --- | --- |
| Linear | 0.563 | 0.340 | 0.438 |
| MLP | 0.580 | 0.344 | 0.420 |
| Deconv | **0.607** | **0.353** | 0.505 |
| Gabor | 0.566 | 0.339 | 0.365 |
| AHP | 0.514 | 0.313 | 0.468 |
| Ours | 0.549 | 0.351 | **0.609** |

Additionally for MNIST images, we compute the accuracy of an MNIST pretrained classifier to measure the quality of the reconstructed digits. The upper bound on the correlation of neuronal responses was 0.944. We skipped testing the neuronal correlation for the “Ours + VAE” reconstructions as they are very similar to the originals and we presume the correlation will be close to the upper bound.

**Table 2.** Evaluation metrics averaged across scans for MNIST images.

| Methods | Pixel-wise correlation | Classification accuracy | Response correlation |
| --- | --- | --- | --- |
| Linear | 0.646 | 0.54 | 0.535 |
| MLP | 0.683 | 0.707 | 0.426 |
| Deconv | 0.783 | 0.947 | 0.730 |
| Gabor | 0.570 | 0.387 | 0.254 |
| AHP | 0.646 | 0.633 | 0.539 |
| Ours | 0.779 | 0.980 | **0.764** |
| Ours + VAE | **0.888** | **0.993** | - |

### A.2 Reconstruction of natural images for all methods

We show reconstructions for some natural images using all different methods (Fig. S1). Hyperparameters for each method are selected as explained in section A.5.

### A.3 Single trial reconstructions

In general, we average the neural activity across many repeats of the image presentation to create reconstructions. Reconstructions on a single presentation of the stimulus are noisier though still qualitatively similar to the original image (Fig. S2). The quality is contingent on the signal-to-noise ratio of the recording apparatus.

**Figure S1.**
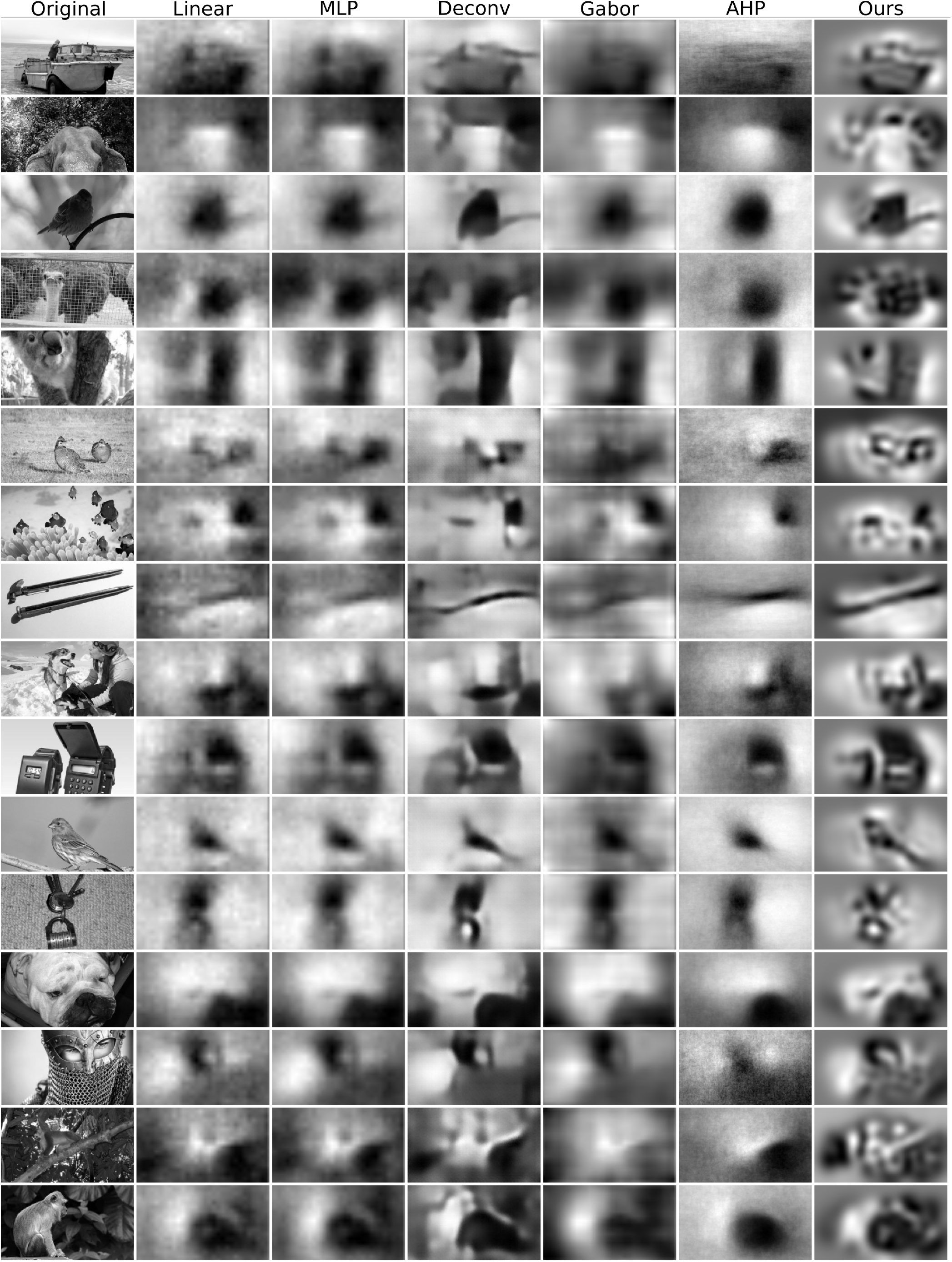
Additional example reconstructions from all methods.

**Figure S2.**
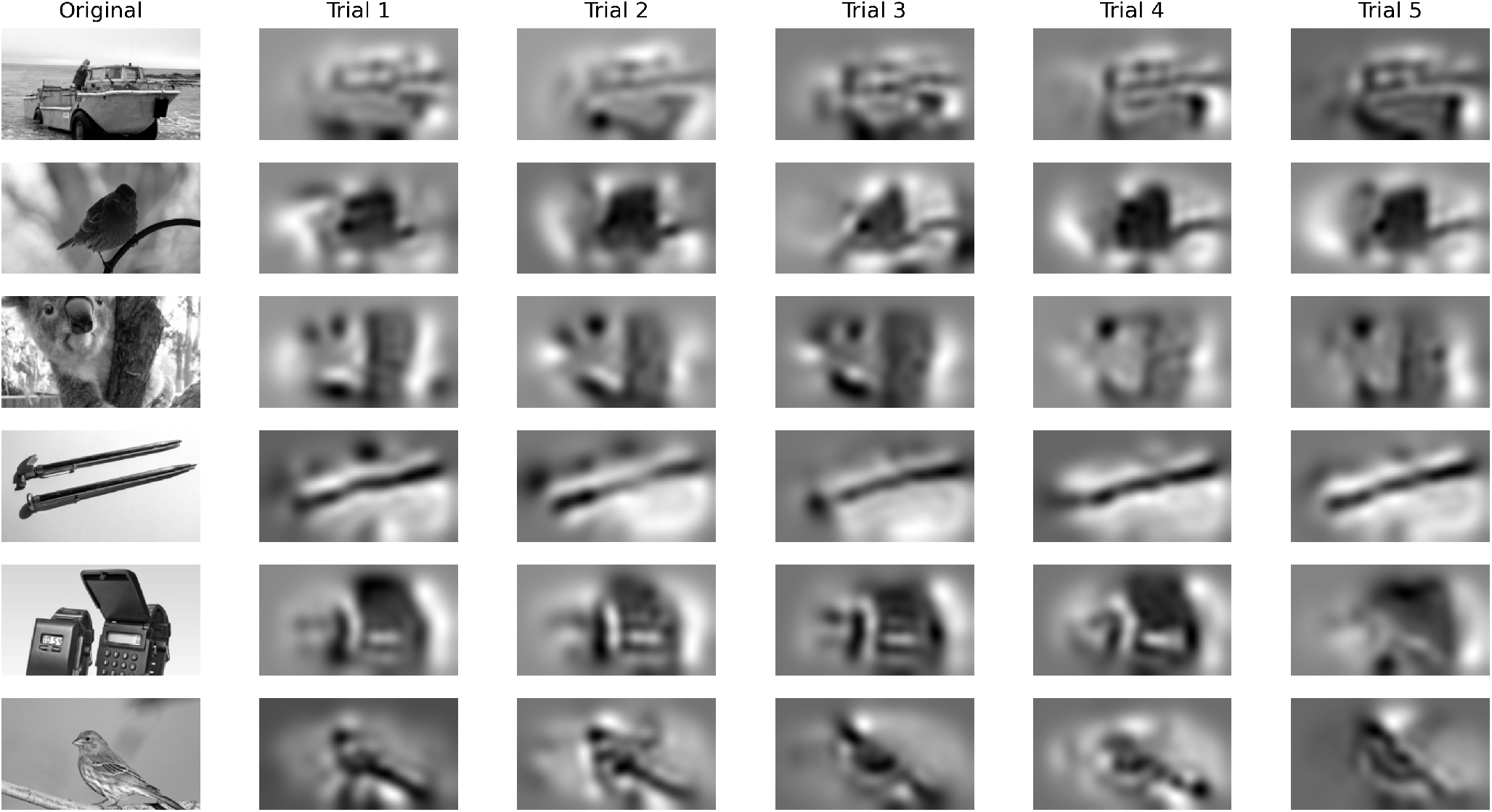
Example reconstructions from a single recording trial. Each column shows the reconstructions computed using the neural responses from a different presentation of the image to the animal. In the main text reconstructions are obtained from responses averaged over many trials.

### A.4 Reconstruction transfer across animals

We tested whether reconstructions from one animal activate the same neurons as the original images in a different animal (Fig. 2g). Reconstructions were generated with recordings from animal 1 and shown alongside the original images to animal 2 in a different session. Recording field-of-view in the second animal is chosen to cover as much of visual cortex as possible. We found that similar trends to those described in the main text hold, namely, that our method still produced the most similar brain responses (0.58 neural response correlation) while the closest next method was Deconv (0.49 correlation). Shuffled shows the correlation of responses from two randomly sampled images, while Orig. shows the correlation of responses to different presentations of the same image (see Sec. A.1); these provide a lower and upper bound for the expected neuronal response correlation given the natural variability of responses (0.27-0.89 correlation).

### A.5 Model selection

For the sake of completeness, we provide the hyperparameters tested for each method. We search a wide range of parameters so each method had a good chance of producing its best results.

- Linear: 4 image sizes ((18, 32), (36, 64), (72, 128), (144, 256)), 11 L2-regularization weights or 7 L1-regularization weight. Total: 72 configurations
- MLP: 4 image sizes ((18, 32), (36, 64), (72, 128), (144, 256)), 5 L2-regularization weights, 2 learning rates (1e-5, 1e-4), 3 hidden layer sizes (1K, 5K, 5K). Total: 120 configurations.
- Deconv: 1 image size (144 × 256), 7 L2-regularization weights, 2 learning rates (1e-4, 1e-3). Total: 14 configurations.
- Gabor: 4 image sizes ((18, 32), (36, 64), (72, 128), (144, 256)), 11 L2-regularization weights or 6 L1-regularization weights, 2 Gabor filter sets (one with 2256 filters as in [5], one with 36992 filters). Total: 136 configurations.
- AHP: 6 number of images to average (1, 3, 10, 32, 136), whether to use a weighted average or not and 2 types of similarity measure between neural responses (correlation or Poisson log-likelihood). Total: 24 configurations.
- Ours: 2 optimization step sizes (100, 500), 9 standard deviations for the gaussian blur ^2^. Total: 18 configurations.

We select the best hyperparameters as those who produce the best results in datasets that do not come from the same animal, i.e., we first find the hyperparameters with the highest average SSIM on the test set in each dataset and then for each dataset, we pick the most common configuration from datasets that do not use the same animal. Although this seems convoluted, we wanted to avoid directly using the test set for each model. Initially, we planned to use an extra validation set per dataset (10% of training images not used for fitting the model) but because these validation images are only shown once to the mice they produce noisier neural responses and using the validation set for model selection produced reconstructions that were unnecessarily smooth (Fig. S3). Structural similarity is already biased towards smother reconstructions and we wanted to make sure we pick a model that still showed detailed reconstruction. The same hyperparameter selection procedure is used for all methods.

**Figure S3.**
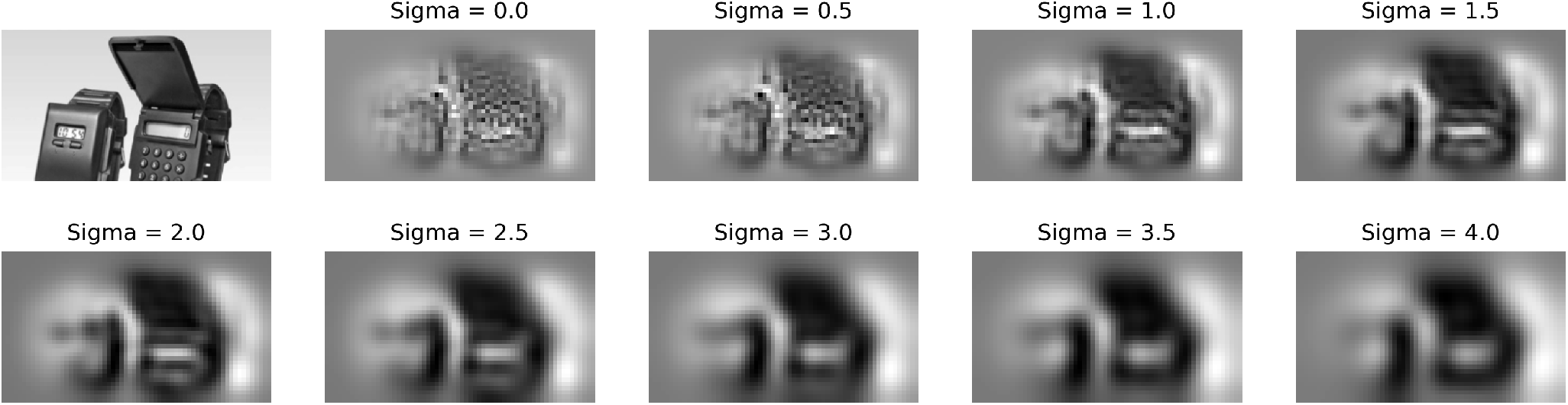
Reconstruction from our method across the range of standard deviations for the gaussian blur. The hyperparameter selection method described above will pick sigma=2.5, using the validation set would have resulted in picking sigma=3.5 which produces overly blurry reconstructions.

### Pixel-wise error

We also computed the reconstruction accuracy (negative of mean squared error) per pixel as a sanity check (Fig. S4). As expected, methods have a similar performance across the entire FOV. Our method only optimizes pixels that are important to elicit the recorded neural responses, thus most pixels outside the field-of-view of the recorded cells are left unchanged, which results in poorer performance (as measured by MSE) near the edges compared to other methods that will try to “outpaint” pixels outside of this FOV using the information from pixels inside the FOV.

**Figure S4.**
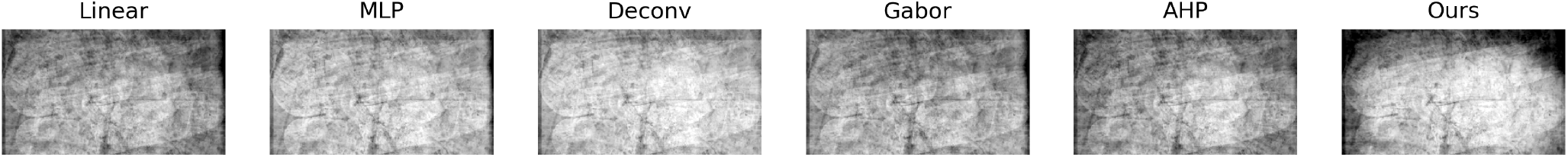
Reconstruction accuracy per pixel (averaged across images) for different methods.

## Footnotes

1 Each scan lasts two hours so we are still able to test all methods in one day.

2 In preliminary experiments, we also tested with adding pixel jittering or using cosine similarity for the neural response comparison but they did not have much effect.

